# Less activity means improved welfare? How pair housing influences pinyon jay (*Gymnorhinus cyanocephalus*) behavior

**DOI:** 10.1101/2024.02.07.579343

**Authors:** London M. Wolff, Jeffrey R. Stevens

## Abstract

The activity level and specific behaviors exhibited by captive animals are crucial indicators of their health and welfare. Stereotypies, or repetitive behaviors that have no apparent function or goal, are performed by animals experiencing poor conditions in their environment and indicate welfare concerns. Changes in the housing environment in particular may have critical influences on behavior and welfare. Here, we measured behavioral changes in a captive pinyon jay (*Gymnorhinus cyanocephalus*) population (n = 12) associated with a shift from single to pair housing. Using automated video processing, we show that pair housing greatly reduced overall activity levels in these birds. The stark reduction in activity was surprising, as we expected that social housing would increase interactions between birds, thus increasing activity levels. Upon further analysis, however, we found that stereotypic behaviors like beak scraping, jumping, pecking, and route tracing decreased after pair housing, whereas a calming behavior—preening— increased. Our results indicate that pair housing may reduce overall activity in pinyon jays; however, this reduction is primarily in stereotypic behaviors.

## Introduction

Bird owners use changes in behavior to track well-being in birds, and a dramatic decrease in activity levels can indicate health problems. But could decreased behavior actually be a sign of lower stress? Currently, activity/movement are offered as proxies for the health and welfare of an animal, with more activity typically linked to improved welfare (Tahamtani et al., 2019; Woods et al., 2022). However, when interpreting reduced activity levels, activity quality or type is rarely considered, highlighting a potential confound: if the behaviors that result in activity are themselves signals of stress, lower activity levels may paradoxically indicate better health. In this paper, we provide evidence of how pair housing of a social bird species is associated with decreased activity, but that the source of this change is decreased stereotypic behaviors, reflecting better, not worse, health and welfare.

Relative to other populations, captive animals are more likely to exhibit stereotypic behaviors, or repetitive behaviors that have no apparent function or goal (Mason, 1991a). Stereotypies, sometimes referred to as abnormal repetitive behaviors, are performed by animals that have in the past or are currently experiencing poor conditions in their environment (Broom, 1983; Mason, 1991a, 1991b; Mellor et al., 2018). Millions of captive birds—whether kept for companionship, education, or poultry production—exhibit stereotypic or other abnormal repetitive behaviors (Mason & Latham, 2004; Mason et al., 2007). These statistics are alarming as these behaviors are known to indicate welfare concerns (Mason, 1991a, 1991b; Rose et al., 2017). In birds, stereotypic behaviors include route tracing, beak scraping, pecking, feather plucking, and repetitive pacing (Garner et al., 2003; Mellor et al., 2018; Woods et al., 2022). Importantly, the presence of stereotypic behavior can tell caretakers that welfare issues are a concern, but caution should be used when making causal assumptions, as there can be a lag between changes in the environment and the reduction of stereotypies if a reduction occurs at all.

Even though stereotypies are not causally interpretable, they typically indicate stress. It is therefore in the best interest of the animal to apply stress-reducing strategies whenever stereotypies appear. Evidence-based solutions can help reduce or eliminate stereotypic behaviors, which is linked to welfare improvements (Mason & Latham, 2004; Williams et al., 2018). The most widely used form of management to combat these abnormal behavior patterns is environmental enrichment (Mason et al., 2007), with a meta-analysis showing enrichment decreasing stereotypic behaviors by 53% (Swaisgood & Shepherdson, 2005). Other possible forms of intervention include punishment, genetic modification, and/or medication. However, these options do not address the underlying issues that cause stereotypies and in some cases can even increase or simply change the type of stereotypy an animal exhibits (Mason et al., 2007). Without addressing the underlying issue of housing or husbandry deficits that cause the stress, reducing stereotypic behaviors themselves is not an ideal endpoint.

Another way to track well-being in an animal population is by observing the number of positive/affiliative behaviors animals exhibit. Naturally occurring behaviors that indicate positive welfare conditions in birds include affiliative behaviors such as allo-preening and food sharing (Clayton & Emery, 2007; Miller et al., in press). This is especially true in highly social species where the connection between pairs of individuals is formed and strengthened through reciprocal preening and the exchange of food (Clayton & Emery, 2007; Duque & Stevens, 2016; Morales Picard et al., 2020). Housing these social species individually may induce stress resulting in stereotypies due to a lack of access to environments that allow the normal and natural functioning of their behaviors (Broom, 1983; Swaisgood & Shepherdson, 2005). Social housing may therefore reduce underlying stress in these species.

## Present Study

Corvids comprise a family of birds found worldwide that includes ravens, crows, magpies, and jays. Due to their sophisticated cognition and varied social structures and feeding ecologies, corvids are a popular study species in the field of animal behavior and cognition (Balda & Kamil, 2002; Clayton & Emery, 2007). With a number of research teams around the world maintaing corvids in captivity to study their behavior, understanding their welfare is critical to this enterprise (Miller et al., in press).

Here, we investigate the effects of housing practices on welfare for pinyon jays (*Gymnorhinus cyanocephalus*), a highly social corvid species that lives in mountainous regions of western North America (Marzluff & Balda, 1992; Balda & Kamil, 1998). Pinyon jays live in flocks ranging from 50 to 500 birds and experience frequent changes in the size and composition of their social groups (Marzluff & Balda, 1992; Wiggins, 2005). They exhibit sophisticated social behaviors, such as inferring transitive social relationships (Paz-y-Mino C et al., 2004), exhibiting social learning (Templeton et al., 1999), food sharing (Duque & Stevens, 2016), prosocial behavior (Duque et al., 2018), and attending to competitor behavior (Vernouillet et al., 2021).

To maintain careful control of food intake and therefore experimental motivation, our colony of pinyon jays has historically been housed individually. However, given the need for social enrichment in corvids (Miller et al., in press), we moved to pair house our birds. We therefore leveraged this change in housing to investigate the effects of different housing conditions on pinyon jay welfare as defined by activity and behavior changes. We hypothesized that moving the birds to larger cages with a conspecific would result in more activity overall due to the new opportunity for social interactions.

Because manually observing and recording behaviors live is so time intensive (Rushen et al., 2012; Whitham & Miller, 2016), we video recorded our pinyon jays before, during, and after they moved to new housing. We then employed an automatic video analysis to quantify activity patterns by tracking pixel changes in the video images. After quantifying their overall activity, we then viewed the video and recorded specific behaviors that the birds exhibited, allowing us to map overall activity onto specific behaviors across the housing transition.

## Materials and Methods

### Subjects

Our study population included 12 (three female) pinyon jays. On moving day, two male birds were replaced with two other males from a different housing room due to unrelated husbandry concerns. As a result, we only focused on the 10 birds that completed all phases of the study when measuring individual behavior.

All birds were wild born, captured in either Arizona or California (United States Fish and Wildlife permit MB694205) between 2006 and 2011. At capture, they were estimated to be between one and three years of age. The birds in this study ranged in age from 10 to 17 years with a mean of 13.1 years. During their time in the lab, all subjects experienced noninvasive cognitive and behavioral experiments and were handled by humans regularly.

The University of Nebraska-Lincoln Institutional Animal Care and Use Committee approved this project (protocol number 2059), and all procedures conformed to the ASAB/ABS Guidelines for the use of animals in research.

### Procedures

#### Housing

Data were collected over a three week period from February 15th, 2021 until March 7th, 2021. During the first week, birds were housed in the single cages that they had been housed in upon entry to the colony (42 × 42 × 60 cm = 0.10 m^3^; Figure 1a). After the first week, we moved each individual animal to their new larger cage with another bird (46 × 96 × 105 cm = 0.46 m^3^; Figure 1b). We label the first week as the *pre-move phase*, the second week as the *during-move phase*, and the third week as the *post-move phase*.

On moving day (February 22nd, 2021), the pinyon jays were placed on either side of the new cage with a divider in place to allow for the animals to acclimate to each other. After an hour of acclimation, we removed the dividers. Lab staff then watched the pairs continuously for the next 20 minutes and periodically for a further two hours to ensure that no animals exhibited aggression or stress. As there was no evidence of negative interactions during this observation period, birds were allowed to remain with their original partners. Of the five pairs created, three were male/female and two were male/male.

#### Recording

We conducted 15-minute video recordings of subjects in their home cage during the three week study period 2-5 times per day (mean 3.7 times per day) between 09:00 and 17:00 CST. In the first week of recording, the animals resided in their original single housing, whereas in the subsequent two weeks, they resided in the new pair housing. Three days prior to the first recording, we habituated the birds to the presence of a tripod and blue colored tape markings on the floor.

For recording sessions, an experimenter placed the camera (GoPro HERO9 Black) on the tripod, turned on the camera, and left the room. No one entered the room during recording sessions. After 15 minutes elapsed, the experimenter re-entered the room, turned off and removed the camera (leaving the tripod), and stored the video recordings. For pair housing, the tripod was adjusted to account for the new height of the paired cages; there were no other changes made to the recording protocol.

### Video processing and analysis

#### Activity levels

To quantify the amount of activity, we used a MATLAB script that calculated the sum of pixel changes across successive frames using the `estimateFlow()` function from the Computer Vision Toolbox. The code started analyzing frames 45 seconds into each video (to eliminate extraneous movement from the birds reacting to the experimenter turning on the camera) and ran until 10 minutes of video had elapsed. Three videos were removed from the analysis due to staff entering the housing room during recording. In total 74 videos with 10 minutes of footage were used in the activity level analysis.

#### Behavior data collection

To further investigate how specific behaviors changed over the three weeks, we coded the birds’ behaviors during week one and three. The first author (LW) created an ethogram of 16 behaviors that were present during the recordings: beak scraping, drinking, feeding, flapping, foraging, head through bars, hopping, jumping, laying down, other, out of view, pecking, perching, playing, preening, route tracing, standing, and walking (see Table 1 for behavior definitions).

**Table 1.** Ethogram of behaviors used to code video.

| Behavior | Definition |
| --- | --- |
| Beak scraping | Bird runs its beak repeatedly back and forth along a branch. |
| Drinking | Bird's head is raised toward water bottle or lowered to the water dish and has beak in contact with the water. |
| Feeding | Bird has its beak inside feeding cup. |
| Flapping | Bird is in an upright position and extends its wings repeatedly. |
| Foraging | Bird pecks or scratches at the ground or manipulates food items. Often includes moving paper in search of food items; however, bird is not only manipulating paper. |
| Head thru Bar | Bird has entire head through the bars of their cage stretching to the outside (can occur within a route-tracing bout). |
| Hopping | Bird jumps up and down on a solid object. Often occurs repeatedly, and it is not using hop to locomote. |
| Jumping | Bird jumps from one perch to another not in a route-tracing bout. |
| Laying down | Bird lays down in an attitude of rest on the floor. |
| Other | Any behavior not belonging to the other categories. |
| Out of View | Bird is not visible for longer than 4 seconds. |
| Pecking | Bird repeatedly pecks at their body (leg band, back, feather, shoulder, etc.). |
| Perching | Bird's feet grasp the perch and bird is not locomoting. |
| Playing | Bird pecks or manipulates an object in the cage other than food or water dish. Unlike foraging it is not directed at searching for food. This can happen while the bird is moving or stationary. |
| Preening | Bird uses its beak to peck, stroke, or comb plumage. |
| Route tracing | Bird follows precise and consistent route within its cage (similar to pacing). |
| Standing | Bird maintains upright position on motionless, extended legs. |
| Walking | Low-speed movement of bird where there legs (not wings) are creating the movement. |

In the post-move phase two birds were present per cage (Figure 1). Because individuals were difficult to identify, it was not possible to tell which of the paired birds were performing a specific behavior. Therefore, we only coded whether a behavior was present in either bird in a cage. To stay consistent across the phases, we also combined both of the birds that would eventually be housed together when coding pre-move phase video data. That is, we combine the behavioral data for each pair throughout both phases. Additionally, our analysis was limited to 10 out of the 12 birds as only 10 birds remained unchanged across phases.

**Figure 1.**
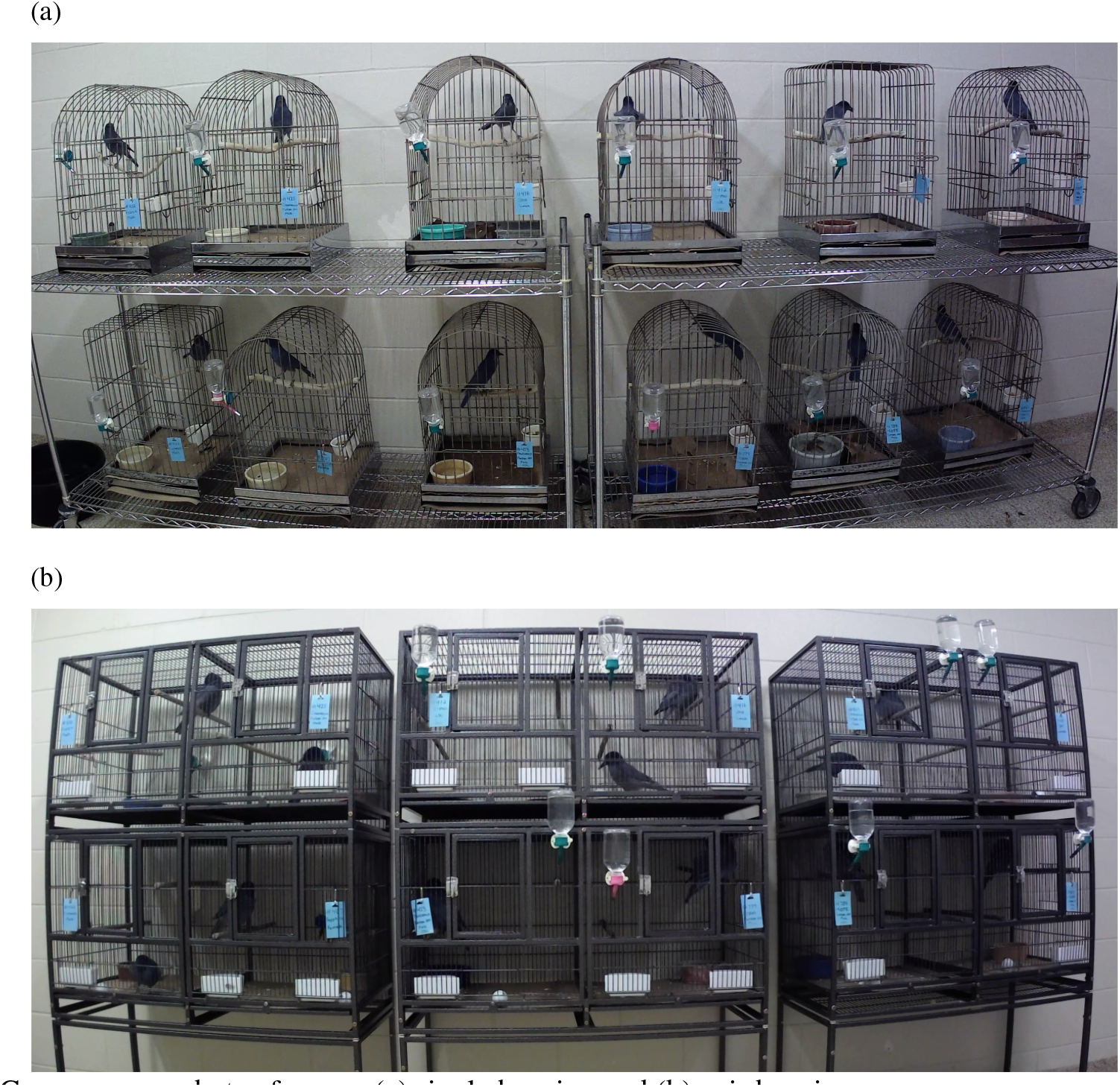
Camera screenshots of cages: (a) single housing and (b) pair housing

For the behavioral analyses, we trimmed the videos to 10 minutes to match the activity level data. We then sampled a 10-second clip per minute per video. The first sample began at the 45-second mark and ended at the 55-second mark. The second sample began at 1 minute 45 seconds and so on, until 9 minutes 45 seconds. We coded 20 recordings from the premove phase and 14 recordings from the post-move phase (we did not code any recordings from the during-move phase). We coded 10 samples per pair per video, resulting in 1700 total samples.

For each of the 16 behaviors on the ethogram, coders recorded the number of times that either bird in a pair exhibited each behavior within every sample. Three coders coded the 1700 samples. To ensure inter-rater reliability, prior to coding the full set, the three coders scored a test set of four videos. LW was aware of the response variable but the other two coders were unware. After training on the ethogram and common issues in coding, each coder received the same randomized subset of four videos to code. We calculated the intraclass correlation of their coded responses using a two-way random effects model for the average of three coders (ICC2k). Based on interpretations from Koo and Li (2016), the intraclass correlation demonstrated good reliability between raters (0.89). To score the full set of videos for analysis, the two unaware coders each scored half of the remaining videos.

### Data analysis

We used R (Version 4.3.2; R Core Team, 2023) and the R-packages *BayesFactor* (Version 0.9.12.4.7; Morey & Rouder, 2024), *easystats* (Version 0.7.0; Lüdecke et al., 2022), *formatstats* (Version 0.0.0.9000; Stevens, 2024), *here* (Version 1.0.1; Müller, 2020), *papaja* (Version 0.1.2; Aust & Barth, 2023), *patchwork* (Version 1.2.0; Pedersen, 2024), *psych* (Version 2.4.1; Revelle, 2024), and *tidyverse* (Version 2.0.0; Wickham et al., 2019) for our analyses. The manuscript was created using *knitr* (Version 1.45, Xie, 2015), *kableExtra* (Version 1.3.4.9000, Zhu, 2023), *rmarkdown* (Version 2.25, Xie et al., 2018), and *papaja* (Version 0.1.2, Aust & Barth, 2023). Data, analysis scripts, supplementary materials, and reproducible research materials are available at the Open Science Framework (https://osf.io/v9r6q/).

Though we present both Bayesian and frequentist statistics (i.e., *p* values), we draw inferences based on Bayes factors because they offer bidirectional information about evidence supporting both the alternative (H_1_) and the null (H_0_) hypotheses. Bayes factors provide the ratio of evidence for H_1_ over evidence for H_0_ (Wagenmakers, 2007; Wagenmakers et al., 2010). Therefore, a Bayes factor of 3 (*BF*_10_=3) indicates three times more evidence for H_1_ than H_0_, whereas a Bayes factor of 1/3 (the reciprocal of 3) indicates 3 times more evidence for H_0_ than H_1_. We interpret Bayes factors based on Wagenmakers et al. (2018), where a *BF*_10_ > 3 is considered sufficient evidence for the alternative hypothesis, *BF*_10_ < 1/3 is considered sufficient evidence for the null hypothesis, and 1/3 < *BF*_10_ < 3 indicate neither hypothesis has evidence supporting it (suggesting the sample size is too small to draw conclusions).

#### Activity levels

We estimated our response variable of activity level by calculating a mean number of pixel changes between video frames. To test the change in activity level over the different phases, we used model selection on linear models calculated with the `lm()` function. We then derived Bayes factors for comparing models from model BIC values using the `test_performance()` function from the *performance* package (Lüdecke et al., 2021). Though we were primarily interested in the effect of phase on activity level, we also included time of day as a potential factor since activity may vary throughout the day. Therefore, we compared four models: (1) an intercept only model `lm(activity ∼ 1),` (2) a phase only model `lm(activity ∼ phase)`, (3) a time of day only model `lm(activity ∼ timeofday)`, and (4) a phase and time of day with no interaction model `lm(activity ∼ phase + timeofday)` (Table 2). We did not include the interaction model in our comparison because we were not interested in how changes in activity level from pre to post-intervention differed depending on the time of day. We calculated Bayes factors comparing each of the models with factors (models 2-4) to the intercept only model (1). We considered the model with the highest Bayes factor as the best fitting model.

#### Behavior data

For behavioral data, we calculated the mean frequency of each behavior per pair for both the pre- and post-move phases. We then conducted frequentist and Bayesian paired t-tests to compare behavior frequency across phases. For the Bayesian t-tests, we employed the `ttestBF()` function from the *BayesFactor* R package (Morey & Rouder, 2024) using default, noninformative priors.

## Results

### Activity levels

Figure 2a shows the range of activity levels across time of day for the three phases. Our comparisons of models (Table 2) showed that the model that included only phase best captured the data. The phase only model had the highest Bayes factor (*BF*_10_ = 2.9×10^23^) compared to the time of day only model (*BF*_10_ = 0.14) and the phase and time of day model (*BF*_10_ = 4.4×10^22^). In fact, there was 6.5 times more evidence favoring the phase only model over the next best (phase and time of day) model. Therefore, phase was an important predictor of activity levels, but time of day was not.

**Table 2.** Model comparison for effect of phase and time of day on activity level.

| Name | Model | AIC | BIC | BF |
| --- | --- | --- | --- | --- |
| Intercept only | activity ~ 1 | 319.0 | 323.6 | NA |
| Phase only | activity ~ phase | 206.3 | 215.6 | $2.9 \times 10^{23}$ |
| Time of day only | activity ~ timeofday | 320.6 | 327.5 | 0.14 |
| Phase and time of day | activity ~ phase + timeofday | 207.8 | 219.3 | $4.4 \times 10^{22}$ |

**Figure 2.**
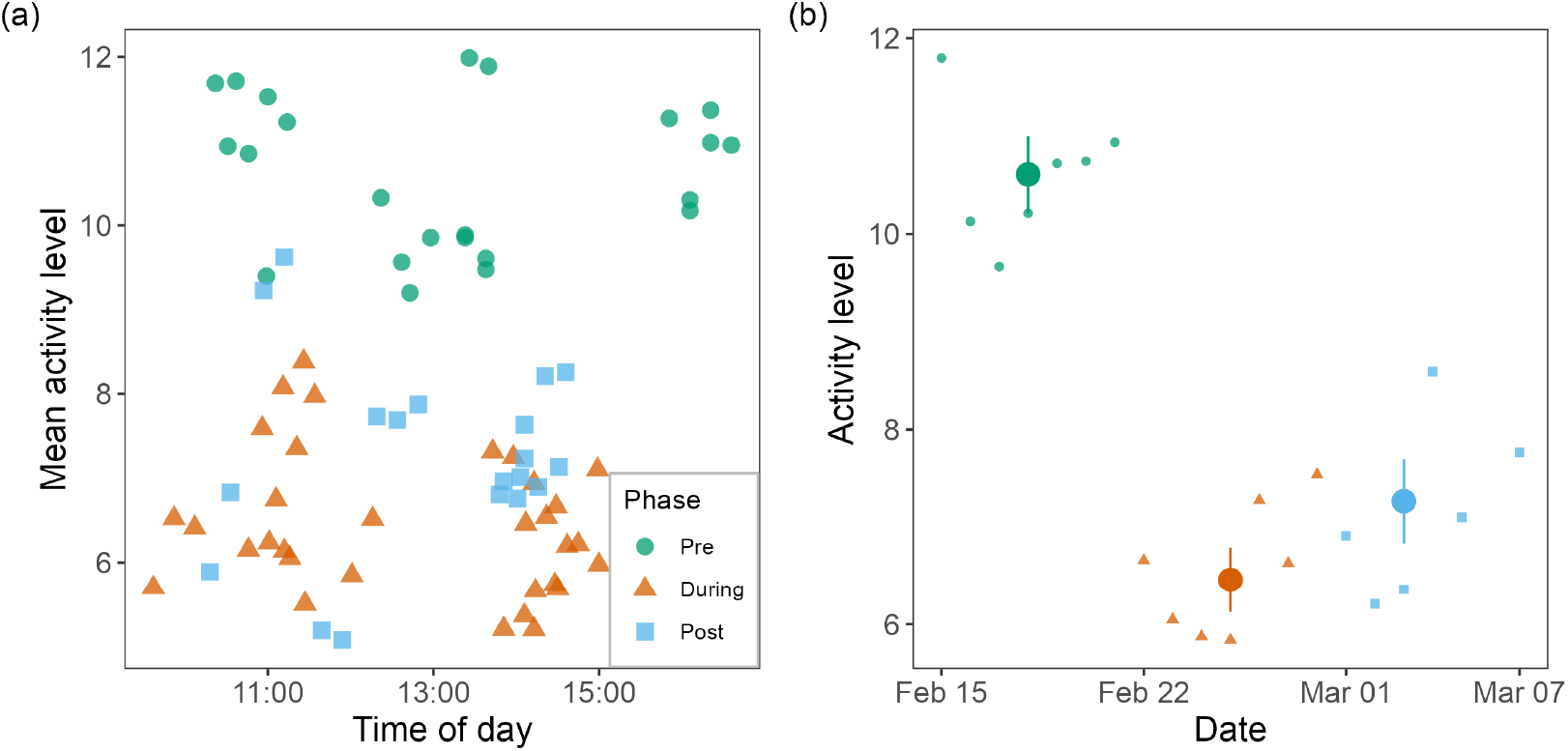
Activity levels. (a) Mean activity levels per sample across time of day for each phase. Points represent mean levels per individual video recording with phase indicated by color and symbol. (b) Mean activity levels per sample across date. Points present mean levels averaged over dates with phase indicated by color and symbol. Dots represent estimated marginal means per phase, and error bars represent 95% confidence intervals.

Since phase was important in predicting activity, we computed pairwise contrasts for the different phases. These contrasts suggest that activity during the pre-move phase was substantially higher than both the during-move phase (Mean difference = 4.15, *t*(71) = 16.1, *p* < 0.001, *d* = 3.8, *BF*_10_ = 3.0×10^20^) and the post-move phase (Mean difference = 3.34, *t*(71) = 11.4, *p* < 0.001, *d* = 2.7, *BF*_10_ = 1.2×10^10^). Further, activity levels increased slightly between the last two phases (Mean difference = -0.81, *t*(71) = -2.9, *p* = 0.004, *d* = -0.7, *BF*_10_ = 7.3). Thus, changing housing greatly reduced overall activity levels (Figure 2b).

### Behavior

The stark reduction in activity was surprising, as we expected that social housing would increase interactions between birds, thus increasing activity levels. After uncovering this finding, we investigated the exploratory hypothesis that reduction in activity was driven by reductions in stereotypic behaviors. Figure 3 shows the mean frequencies for all of the behaviors, along with Bayes factors and p-values for the paired t-tests comparing frequencies in the pre- and post-move phases. Of the 16 behaviors, we observed a decrease in beak scraping, feeding, foraging, jumping, pecking, playing, route tracing, and walking. We observed an increase in perching and preening. We did not have enough evidence to detect differences or lack of differences in drinking, flapping, head thru bar, hopping, laying down, or standing.

**Figure 3.**
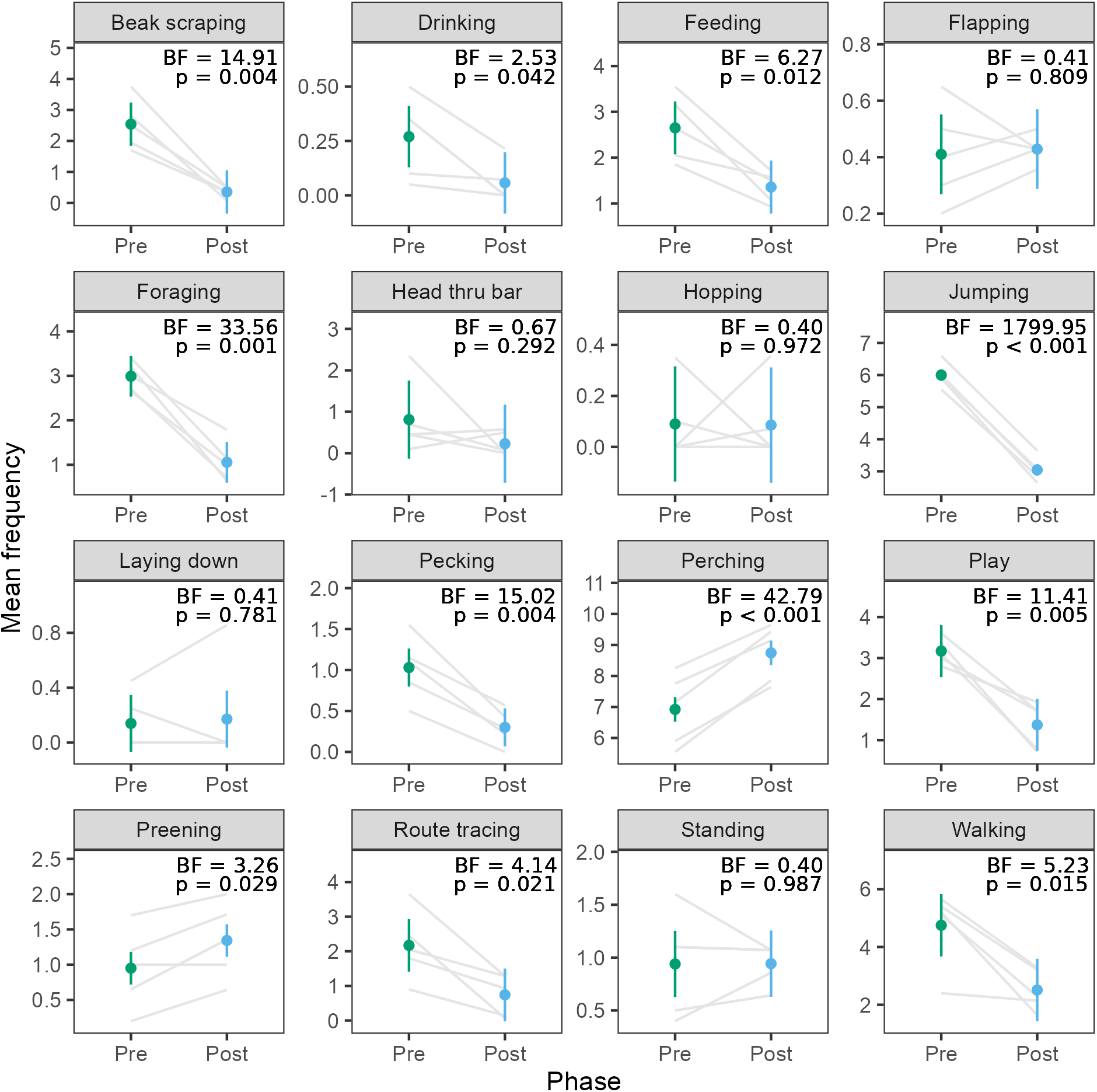
Mean frequencies of 16 behaviors in the pre- and post-move phases. Grey lines connect means for each of the five bird pairs. Dots represent overall means per phase, and error bars represent within-pair 95% confidence intervals.

## Discussion

We examined behavioral changes in pinyon jays during two husbandry interventions of a larger cage and pair housing. After the cage change, birds decreased their activity levels as measured by overall pixel changes during video recording. This dramatic drop in activity was surprising and motivated a more extensive follow-up analysis examining the frequencies of specific behaviors. This exploratory analysis indicated that perching and preening increased in frequency after the cage change, while beak scraping, feeding, foraging, jumping, pecking, playing, route tracing, and walking decreased after the cage change. Thus, moving to pair housing substantially altered behavior in the pinyon jays.

### Animal Welfare Implications

This study highlights a crucial distinction in the assessment of captive animal welfare: less activity does not necessarily imply poor welfare or increased stress. Rather, it is one facet of animal behavior that must be examined when determining animal welfare. Our data in particular show that moving from single to pair housing can result in an overall reduction in activity. Yet that reduction does not occur uniformly in all behaviors. Our birds demonstrated reductions in stereotypic behaviors associated with stress such as beak scraping, jumping, pecking, and route tracing. Therefore, the pair housing seems to have reduced this repetitive behaviors. However, it also decreased seemingly positive behaviors such as foraging, playing, and walking. These behaviors might have decreased because the social enrichment associated with pair housing substituted for other forms of physical enrichment that the birds engaged in to maintain their own psychological welfare. Having a social partner present may have replaced the need to engage in these other activities. We also observed an increase in preening and perching. These behaviors may indicate a reduction in stress, where the animals feel comfortable enough in their environment that they can rest calmly and engage in self-care. Thus, overall, the move from single to pair housing seems to have reduced stress-related behaviors and increased calming behaviors.

The growing prevalence of automated behavior assessment systems such as video recording, accelerometers, and GPS devices can facilitate the large-scale collection of activity data (Rushen et al., 2012; Whitham & Miller, 2016). However, researchers and animal caretakers must be mindful that overall patterns of activity do not necessarily provide a complete assessment of welfare. Measuring specific behaviors associated with stress and calm are critical to assessing welfare and formulating care plans. It is imperative to recognize that when employing activity measures as an indicator of health in captive animals, the absence of certain behaviors is not inherently problematic. Automated processes can be useful in assessing animal welfare, but human observers provide an invaluable perspective on the health and well-being of captive animals.

### Limitations

Though our data provide intriguing insights into the effects of housing changes on captive bird welfare, we note several limitations of our study. First, this study involves a relatively small population of 12 birds. Of course, individual differences are a critical component of animal behavior and welfare (Stamps et al., 2012; Dingemanse & Wolf, 2013; Richter & Hintze, 2019). Interestingly, though some of the behaviors that we scored showed quite a bit of variability, others were quite consistent. Beak scraping, foraging, jumping, and play all showed both consistent frequencies before the housing change and consistent drops in frequencies after the change. Other behaviors such as foraging, pecking, perching, route tracing, and walking showed variability in the initial frequencies but consistent decreases (or increases) after the housing change. Thus, despite a relatively small sample size, most of our behavioral measures show consistent patterns across individuals. Moreover, the logistics of viewing videos of birds in pair housing did not allow us to identify and attribute behaviors to specific birds. Instead behaviors were coded across bird pairs. Our findings are therefore limited to generalizations, not claims about specific individual reduction or increases in individual behaviors. Larger samples with individually identifiable subjects would provide more confidence about the generalizability of results.

A second limitation is the advanced age of our birds (10 to 17 years old). Younger birds have different abilities to rebound to new and novel changes in their environments (Greenberg, 2003). It is possible that the reduction in activity and behaviors in our birds could have been an adverse reaction to the changing in housing. The lack of movement and increased perching could indicate more of a ‘freezing’ response to the stress of the change. While this is possible, the increase in preening indicates more comfort with their surroundings. However, replicating this work with a larger sample size, a more diverse age range of birds, and perhaps more physiological measures of stress (e.g., cortisol, heart rate levels) could clarify the effects of pair housing on bird welfare.

Finally, we only recorded behavior for two weeks after the housing change. Though it was a small difference, activity levels in the third week increased over the second week. It is possible that the activity levels would have continued to increase over time. Therefore, we cannot claim that the behavioral differences observed here represent a sustained or permanent change in behavior. Rather we can only offer a snapshot in time that needs longer-term studies to determine if these activity patterns stay consistent as the pairs become more acquainted.

## Conclusion

This research investigated how pinyon jays showed paradoxically lower activity levels after moving from single to pair housing. Upon further video analysis we found that the stereotypic behaviors of beak scraping, jumping, pecking, and route tracing decreased after pair housing, whereas a calming behavior—preening—increased. Our findings suggest that pairing pinyon jays may decrease their overall activity, but this decrease is mainly observed in stereotypical behaviors. Further research is needed to see if this reduction in activity is sustained over time following initiation of pair housing.

## Acknowledgements

This research was funded by a U.S. National Science Foundation grant (NSF-1658837). We would like to thank our amazing lab managers Kylie Hughes and Katie Carey for collecting data and helping the lab run smoothly and our research assistants Toria Biancalana, Bailey Wilson, and Hailey Wilson for collecting data and Rachel Bruner and Isaac Martinez for helping code the behavioral video data. We are grateful to Tierney Lorenz for comments on an early version of the manuscript.

## Author Contributions

**Wolff:** Conceptualization, Methodology, Validation, Investigation, Data Curation, Writing – Original Draft, Writing-review & editing, Supervision, Project administration.

**Stevens:** Conceptualization, Methodology, Software, Formal Analysis, Resources, Data Curation, Writing – Original Draft, Writing-review & editing, Visualization, Supervision, Funding acquisition.

## Data Availability

The data and analysis code are available at: https://osf.io/v9r6q/.

## Ethics Approval

All procedures were conducted in an ethical and responsible manner, in full compliance with all relevant codes of experimentation and legislation and were approved by the UNL Institutional Animal Care and Use Committee (protocol #2059).

